# Investigating the effects of age and hearing loss on speech intelligibility and amplitude modulation frequency selectivity

**DOI:** 10.1101/2023.07.20.549841

**Authors:** Jonathan Regev, Johannes Zaar, Helia Relaño-Iborra, Torsten Dau

## Abstract

The perception of amplitude modulation (AM), characterized by a frequency-selective process in the modulation domain, is considered critical for speech intelligibility. Previous literature has provided evidence of an age-related decline in AM frequency selectivity. Additionally, a notable sharpening of AM tuning with hearing loss has been observed, which was proposed to be related to a perceptual advantage resulting from peripheral compression loss. This study explores whether such changes in AM tuning with age and hearing loss contribute to the speech intelligibility challenges that older listeners often face in noisy environments. Young (n=10, 22-28 years) and older (n=9, 57-77 years) listeners with normal hearing as well as older listeners with hearing impairment (n=9, 64-77 years) participated in the study. All had previously taken part in studies on AM tuning. Speech-reception thresholds (SRTs) were collected in conditions including stationary, fluctuating, and competing-speech maskers. The results revealed an age-related increase in SRTs, with an additional negative impact of hearing loss. Beyond age and audibility, a measure of AM tuning significantly contributed to explaining the variance in SRTs across listeners for specific maskers. These findings motivate further exploration of the relationship between AM frequency selectivity and speech intelligibility in noise.

## I. INTRODUCTION

Temporal envelope fluctuations (also known as “amplitude modulation” or AM) refer to the slow variations in overall amplitude that characterize most sounds in daily life (e.g., Singh and Theunissen, 2003; Elliott and Theunissen, 2009). The auditory system exhibits remarkable sensitivity to AM, as demonstrated by physiological studies (see Joris *et al*., 2004 for a review) and behavioral investigations of AM detection (e.g., Viemeister, 1979; Kohlrausch *et al*., 2000). Similar to masking effects in the audio-frequency domain, the detectability of a target AM is disrupted by the presence of distinct masking AM components. AM masking has been extensively studied in behavioral investigations (e.g., Bacon and Grantham, 1989; Houtgast, 1989; Dau *et al*., 1997; Verhey *et al*., 2003) as well as through recent neurophysiological research (Viswanathan *et al*., 2021). Behavioral studies of AM masking have provided evidence of a frequency-selective process in the modulation domain, often conceptualized as a series of bandpass modulation filters known as the modulation filterbank (e.g., Dau *et al*., 1997, 1999; Ewert and Dau, 2000; Ewert *et al*., 2002). Computational modelling studies have shown that this concept significantly contributes to quantitatively predicting experimental data from various psychoacoustic tasks (e.g., Dau *et al*., 1997, 1999; Jepsen *et al*., 2008; Jepsen and Dau, 2011; McDermott and Simoncelli, 2011; McWalter and Dau, 2017), underscoring the importance of AM in auditory processing and perception.

Beyond its role in ‘simple’ listening tasks, AM has been argued to be crucial for accurate speech intelligibility (Houtgast and Steeneken, 1985; Drullman *et al*., 1994; Smith *et al*., 2002; Elliott and Theunissen, 2009). The significance of AM in speech was highlighted by vocoding studies, which demonstrated that intact envelope information conveyed through a few broad spectral bands was sufficient for precise speech recognition in quiet, even when spectral cues were greatly reduced (e.g., Shannon *et al*., 1995; Dorman *et al*., 1997; Xu *et al*., 2005). Based on this, AM masking has been proposed as the primary mechanism underlying speech intelligibility in noise, whereby speech-reception thresholds (SRTs) - reflecting the signal-to-noise ratio required for 50% speech recognition - are related to the modulation-domain signal-to-noise ratio (SNR_env_) between the speech and masker (e.g., Dubbelboer and Houtgast, 2008; Jørgensen and Dau, 2011).

The crucial role of AM masking in speech intelligibility in noise was also emphasized by studies using maskers designed to contain AM components or to have maximally flat envelopes, which suggested that so-called stationary maskers, usually associated with energetic masking (EM) in the audio-frequency domain, effectively function as AM maskers due to their inherent envelope fluctuations (Stone *et al*., 2012; Stone and Moore, 2014). Computational auditory modeling studies have successfully incorporated these concepts by employing the SNR_env_ (Jørgensen and Dau, 2011; Jørgensen *et al*., 2013; Chabot-Leclerc *et al*., 2014) or correlation-based measures (Relaño-Iborra *et al*., 2016, 2019; Steinmetzger *et al*., 2019) at the output of the modulation filterbank as decision metrics. Based on these modeling studies, the spectral decomposition of the stimulus envelope carried out by the modulation filterbank, i.e., AM frequency selectivity, has been suggested as an important factor for accurately predicting SRTs (see Relaño-Iborra and Dau, 2022 for a review of different speech intelligibility models using modulation filtering).

Regev *et al*. (2023, 2024a) investigated the effects of age and hearing loss on the selectivity of hypothetical modulation filters using a perceptual AM masking task. Regev *et al*. (2023) found a general reduction in AM frequency selectivity associated with age, resulting in broader masking patterns among older compared to young listeners with normal hearing (referred to as hereafter as NH listeners). Conversely, Regev *et al*. (2024a) demonstrated a general sharpening of masking patterns among young and older listeners with hearing impairment (referred to hereafter as HI listeners) compared to NH listeners of similar age. This finding suggested that sensorineural hearing loss was accompanied by a perceptual benefit that partially offset the detrimental effects of age, possibly due to the HI listeners’ ability to exploit an increased internal representation of AM depth resulting from reduced peripheral compression when “listening in the dips” of the masker. Hence, Regev *et al*. (2024a) proposed that the sharpening of the masking patterns found in HI listeners was an “apparent” effect, reflecting the loss of peripheral compression rather than an actual greater selectivity of the hypothetical modulation filters. In other words, Regev *et al*. (2024a) proposed that older HI listeners also experience an age-related reduction in AM frequency selectivity, which is concealed by a perceptual benefit stemming from the loss of peripheral compression. Such age-related changes in the sharpness of the hypothetical modulation filters may affect speech intelligibility in noise by altering the amount of AM masking at the output of the modulation filters.

The effects of age and hearing loss on speech intelligibility have received significant attention in the literature. Hearing loss is known to cause a decline in speech intelligibility in both quiet and noisy environments (e.g., Glasberg and Moore, 1989; Humes and Christopherson, 1991; Frisina and Frisina, 1997; see Humes and Dubno, 2010 for a review). In addition to audibility loss, speech intelligibility challenges associated with hearing loss have been linked to various supra-threshold deficits, including reduced frequency selectivity, poorer temporal fine structure processing, reduced recovery from forward masking, poorer AM depth discrimination, and loudness recruitment (Glasberg and Moore, 1986, 1989; Oxenham and Moore, 1997; Moore and Oxenham, 1998; Moore, 2003; Lorenzi *et al*., 2006; Strelcyk and Dau, 2009; Schlittenlacher and Moore, 2016; Gallun *et al*., 2022). Aided speech intelligibility performance has also been shown to be strongly correlated with deficits in spectro-temporal modulation detection in HI listeners (e.g., Bernstein *et al*., 2016; Zaar *et al*., 2023, 2024). Recent neurophysiological studies suggested that poorer speech intelligibility in HI listeners may be related to increased neural coding of the envelope (Millman *et al*., 2017; Goossens *et al*., 2018; Decruy *et al*., 2020). This effect may be partly attributed to an increased internal representation of the AM depth resulting from the loss of cochlear compression commonly associated with sensorineural hearing loss (Moore *et al*., 1996; Jennings *et al*., 2018). In addition to the effects of hearing loss itself, age and reduced cognitive function have been found to contribute to the decline in speech intelligibility observed in older HI listeners (e.g., Dubno *et al*., 1984; Humes and Christopherson, 1991; Frisina and Frisina, 1997; Humes, 2002, 2007; Humes *et al*., 2006). This suggests that cognitive and age-related perception deficits may also play a role in speech intelligibility challenges.

To investigate these potential mechanisms, cross-sectional studies focusing on age-related effects have been conducted, often involving listeners with no or mild hearing loss. These studies have consistently shown a detrimental effect of age on speech intelligibility in the presence of competing speech maskers (Rajan and Cainer, 2008; Helfer and Vargo, 2009; Schoof and Rosen, 2014; Füllgrabe *et al*., 2015; Gordon-Salant and Cole, 2016; Goossens *et al*., 2017; Decruy *et al*., 2019; Sobon *et al*., 2019). However, the effects of age in stationary and fluctuating noise have been less consistent. Some studies found differences between young and older listeners (e.g., Goossens *et al*., 2017; Decruy *et al*., 2019; Sobon *et al*., 2019), while others did not (Rajan and Cainer, 2008; Helfer and Vargo, 2009; Schoof and Rosen, 2014). Overall, the age-related increase in SRTs has been more pronounced for speech maskers than for stationary or fluctuating noises (e.g., Goossens *et al*., 2017; Decruy *et al*., 2019; Sobon *et al*., 2019). The observed age-related decline in speech intelligibility has been suggested to be mediated by deficits in temporal processing (e.g., Gordon-Salant *et al*., 2006; Anderson *et al*., 2012; Füllgrabe *et al*., 2015; Goossens *et al*., 2016; Decruy *et al*., 2019; McFarlane and Sanchez, 2024) and by reduced cognitive abilities (Humes *et al*., 2012; Füllgrabe *et al*., 2015; Gordon-Salant and Cole, 2016).

Since AM frequency selectivity has been identified as a significant factor contributing to accurate speech intelligibility, it is reasonable to suggest that the age-related decline in NH listeners’ AM selectivity reported by Regev *et al*. (2023) may contribute to the aforementioned challenges in speech intelligibility. A reduction in AM tuning would likely result in increased AM masking, thus disrupting the intelligibility of the target speech. Similarly, it is hypothesized that older HI listeners also experience an age-related reduction in AM selectivity that contributes to a decline in speech intelligibility. Although the increased AM frequency selectivity observed in older HI listeners compared to older NH listeners (Regev *et al*., 2024a) might suggest reduced AM masking with hearing loss, it was proposed that this greater AM selectivity in HI listeners is an apparent effect caused by reduced cochlear compression, rather than a true sharpening of the hypothetical modulation filter. Consequently, the contribution of an age-related reduction in AM selectivity to speech intelligibility difficulties in older HI listeners should become evident once hearing loss is accounted for. Furthermore, reduced cochlear compression is associated with supra-threshold deficits that impair speech intelligibility (e.g., Glasberg and Moore, 1989; Oxenham and Moore, 1997; Moore and Oxenham, 1998; Moore, 2003). This aligns with studies showing that HI listeners exhibit higher SRTs than age-matched NH listeners even when audibility is restored (Goossens *et al*., 2017).

The present study investigated whether the results of speech intelligibility tests support the assumption that an age-related decline in AM frequency selectivity negatively impacts speech intelligibility in both older NH and older HI listeners, once the effects of cochlear compression loss in HI listeners are accounted for. SRTs were collected for speech in noise, building upon the previous studies on AM frequency selectivity (Regev *et al*., 2023, 2024a), using a subset of the participants. Young NH, older NH, and older HI listeners were tested on sentence recognition in various maskers, including speech-shaped noise, fluctuating noise, and competing talkers. It was hypothesized that age-related changes in AM frequency selectivity would be linked to increases in SRTs for the older NH and HI listeners.

## II. METHODS

### A. Listeners and previously published data

All young NH and older NH listeners tested by Regev *et al*. (2023) as well as all older HI listeners tested by Regev *et al*. (2024a) were invited to participate in the present study. Ten young NH listeners, aged 22-28 years (mean age = 24.7 years, eight female), nine older NH listeners, aged 57–77 years (mean age = 66.9 years, six female), and nine older HI listeners, aged 64-77 years (mean age = 71.2 years, 2 female), accepted this invitation. All listeners were native Danish speakers. The experiment was conducted monaurally in the same test ear as used by Regev *et al*. (2023, 2024a). Figure 1 shows the audiograms in the participants’ test ears. All young NH listeners had audiometric pure-tone thresholds (PTTs) ≤ 20 dB HL at frequencies ranging from 125 Hz to 8 kHz. All older NH listeners had PTTs ≤ 20 dB HL at frequencies ranging from 125 Hz to 4 kHz, PTTs ≤ 25 dB HL at 6 kHz, and PTTs ≤ 45 dB HL at 8 kHz. The older HI listeners exhibited a high-frequency sloping hearing loss, classified as an N2 or N3 standard audiogram (Bisgaard *et al*., 2010),^1^ and audiometric thresholds in the range of 50-60 dB HL at 3 kHz as required by Regev *et al*. (2024a). All HI listeners had sensorineural hearing loss, with air-bone gaps ≤ 15 dB at 1, 2, 3, and 4 kHz.^2^

**Figure 1.**
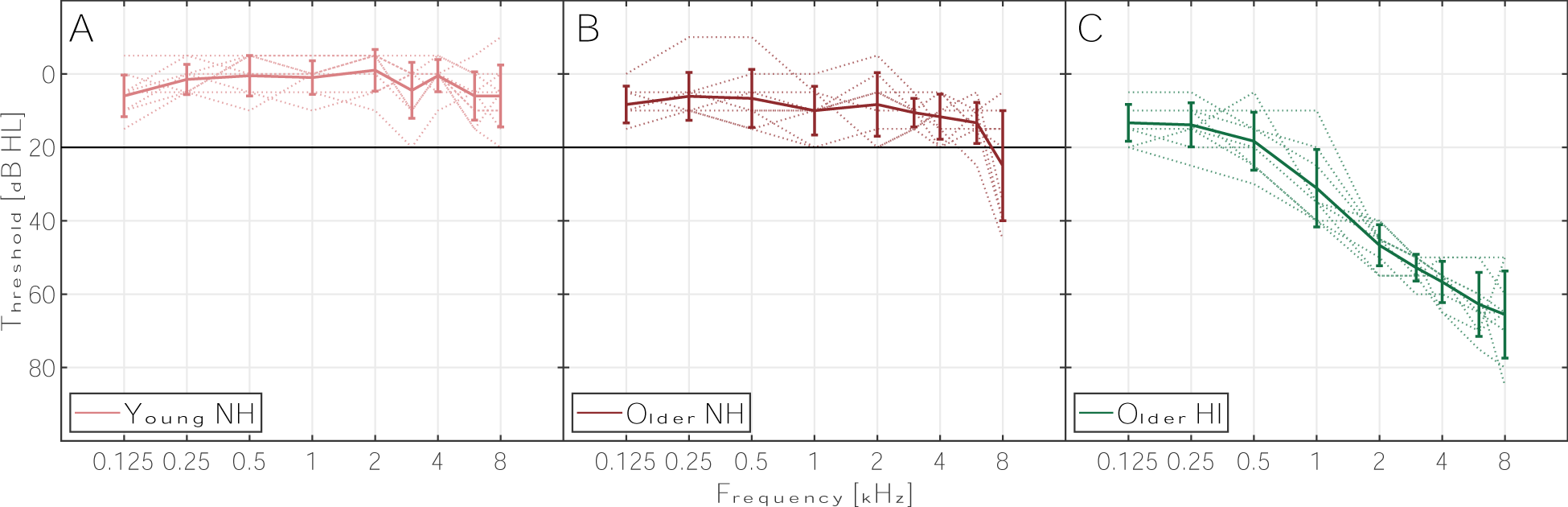
(Color online): Individual and average audiograms of the test ear for each listener group. The thin dashed lines show the individual audiograms, while the dark solid lines represent the average and standard deviation across listeners. Panel A shows the audiograms for the young NH listeners, with the black horizontal line indicating the 20 dB HL normal-hearing criterion. Panel B displays the audiograms for the older NH listeners, also with the black horizontal line denoting the 20 dB HL normal-hearing criterion. Panel C presents the audiograms for the older HI listeners.

Prior to this study, all listeners took part in the AM frequency selectivity studies conducted by Regev *et al*. (2023, 2024a). The corresponding datasets are publicly available from Regev *et al*. (2024c, 2024b). In those studies, masked-threshold patterns (MTPs) were used to assess the masked detection threshold of a target AM across different masker center frequencies, providing an indication of AM frequency selectivity. Figure 2 indicates an example of the MTPs for a 4-Hz target modulation frequency, where the age-related reduction in AM selectivity is evident on both skirts and the sharpening of the MTP with hearing loss is evident on the low-frequency skirt (see Regev *et al*., 2023, and 2024a for a detailed discussion of the possible origins of these effects). A measure of AM frequency selectivity at the individual level can be derived from the MTPs by computing the average range spanned by the MTP (i.e., the MTP’s average dynamic range; DR_MTP_), where a larger dynamic range is indicative of greater AM selectivity. This is done by identifying the highest masked threshold as the peak of the MTP, computing the difference between the peak and the lowest masked threshold on each side of the peak, and averaging the resulting dynamic range across both sides. If the peak of the MTP is on either extreme, the DR_MTP_ is taken as the dynamic range across the whole MTP. An illustration of this derivation is provided in supplementary Figure 1. This measure was derived for the 4-Hz MTP of each listener (with the exception of one older HI listener for whom a MTP at 4 Hz could not be obtained by Regev *et al*., 2024a), since the 4-Hz modulation frequency has been suggested to be particularly important for speech perception (e.g., Drullman *et al*., 1994; Arai *et al*., 1999; Greenberg *et al*., 2003; Elliott and Theunissen, 2009; Varnet *et al*., 2017) and showed the largest effect sizes across listener groups in Regev *et al*. (2023, 2024a). The resulting individual quantifications of AM frequency selectivity are shown in Figure 3 for each listener group. As reported by Regev *et al*. (2023, 2024a), the MTPs and corresponding dynamic ranges indicate a general decline in AM frequency selectivity with age, which is partially “compensated” by the presence of hearing loss.

**Figure 2.**
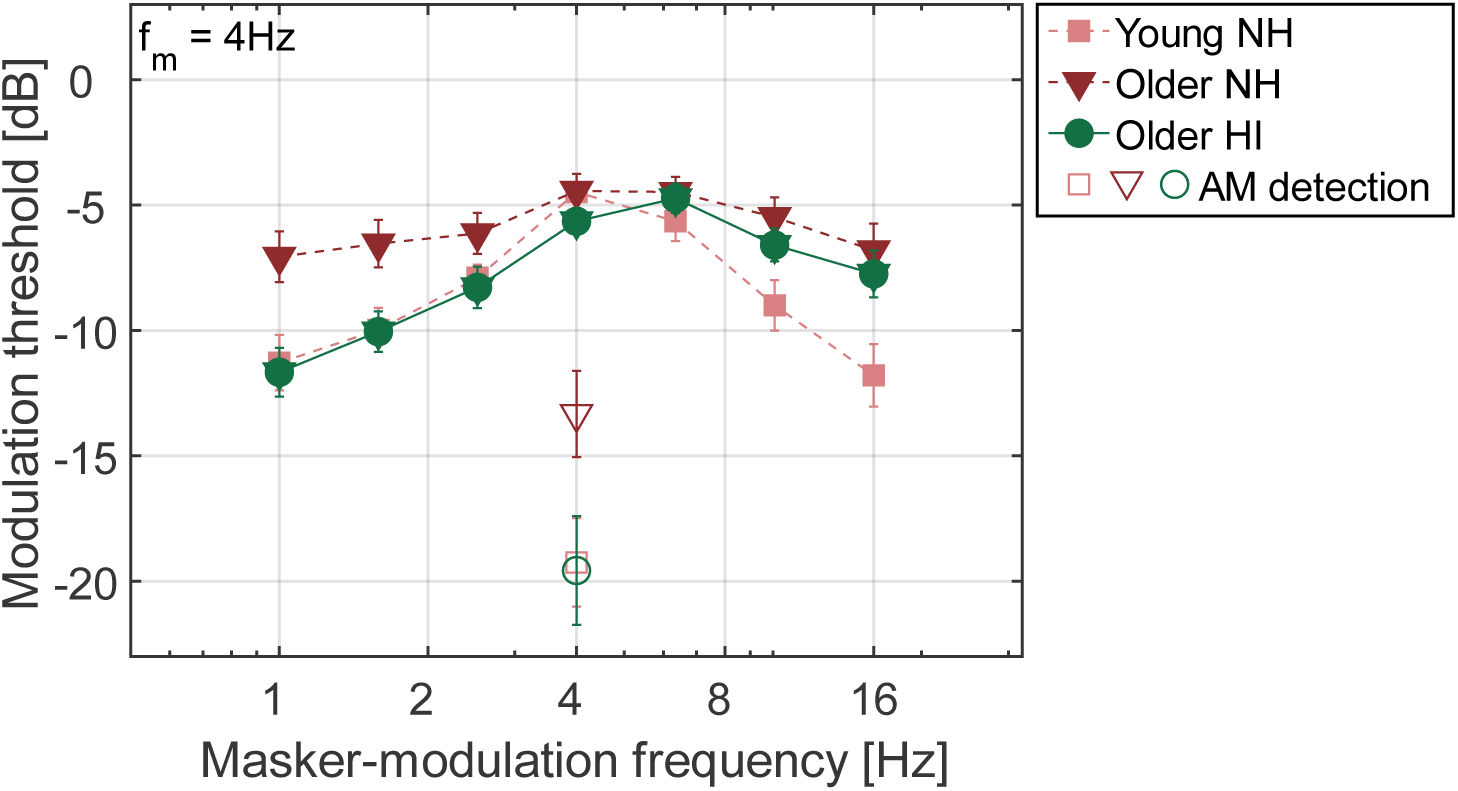
(Color online): Average MTPs and AM detection thresholds for a 4-Hz target modulation frequency. The results for the young NH group are presented as dashed lines and squares, the older NH group as dashed lines and triangles, and the older HI group as continuous lines and circles. The bottom open symbols represent the average AM detection threshold for each group. The means and standard errors are shown. Data adapted from Regev *et al*. (2023, 2024a).

**Figure 3.**
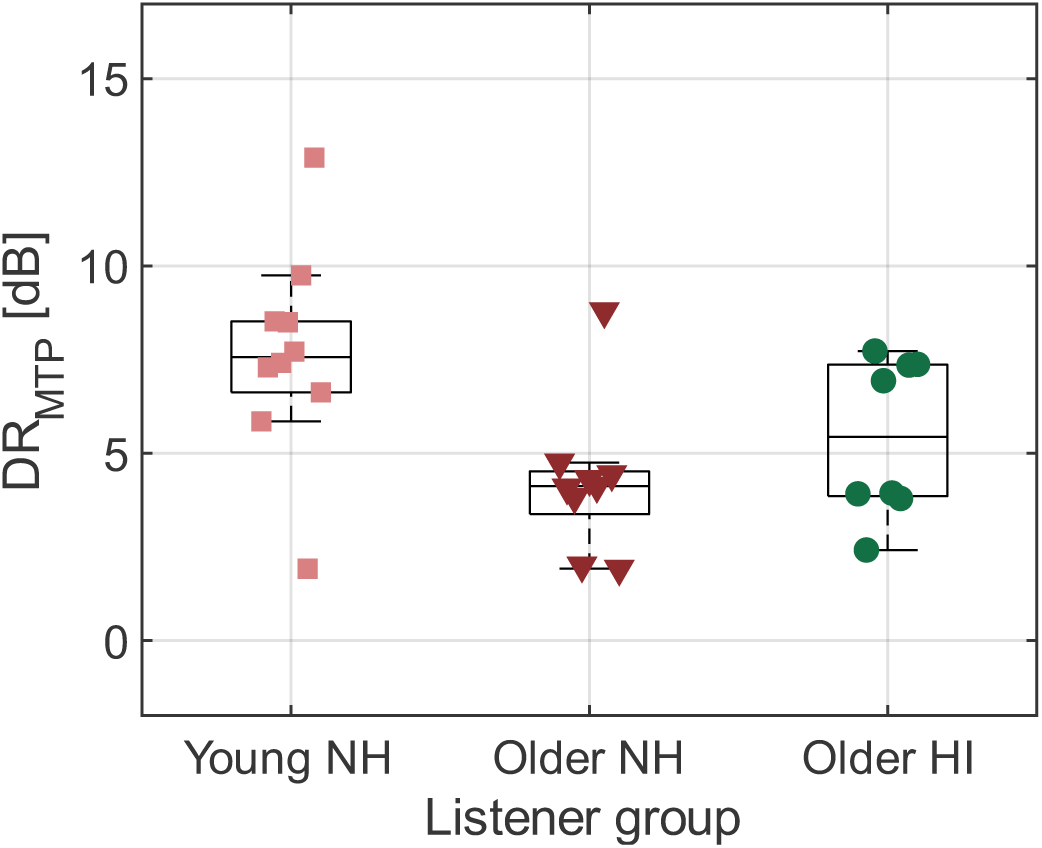
(Color online): Boxplots displaying the average dynamic range of the 4-Hz MTP (DR_MTP_) for all listeners. The data for the young NH listeners are represented by squares, older NH by triangles, and older HI by circles. Data derived from Regev *et al*. (2023, 2024a).

Regev *et al*. (2023, 2024a) also evaluated working-memory capacity using a reverse digit span (RDS) test, which was administered binaurally following the procedure described in Fuglsang *et al*. (2020). For the HI listeners, a linear gain was applied based on the Cambridge formula (CamEq; Moore and Glasberg, 1998), with a maximum gain of 30 dB at any frequency. The older NH listeners exhibited significantly lower average RDS scores (0.40 on a normalized scale) compared to the young NH and older HI listeners (0.60 and 0.52, respectively).^3^

All participants provided written informed consent and were offered financial compensation for their participation. The study received ethical approval from the Science Ethics Committee of the Capital Region of Denmark (reference H-16036391).

### B. Apparatus

The listening test was conducted in a single-walled soundproof booth. The experiment was conducted using MATLAB (The Mathworks Inc., 2015). The stimuli were digitally played using SoundMexPro (HörTech, Oldenburg, Germany), converted to analog using an RME Fireface soundcard (Haimhausen, Germany), and played monaurally through Sennheiser HDA200 headphones (Wedemark, Germany). The output level was calibrated using a GRAS RA0039 ear simulator (Holte, Denmark) and the frequency response of the headphones was compensated using an inverse digital filter to achieve a flat response at the eardrum. The listeners received no feedback during the test.

### C. Procedure and stimuli

Speech intelligibility in noise was assessed using the Danish Hearing in Noise Test (HINT; Nielsen and Dau, 2011). The HINT speech material consists of 10 test lists and 3 training lists, each containing 20 five-word sentences. The sentences were spoken by a male speaker with an estimated average fundamental frequency (*F_0_*) of 123 Hz. The speech signals were combined with five different masking noises: i) a stationary speech-shaped noise (SSN); ii) a fluctuating ICRA-5 noise (Dreschler *et al*., 2001) resembling a single male talker in terms of spectral and temporal characteristics; iii) a single male competing talker reading a news piece (*F_0_* = 118 Hz); iv) a single female competing talker reading a news piece (*F_0_* = 173 Hz); and v) a ‘cocktail party’ condition. The recordings of the competing talkers had previously been edited to reduce long pauses to 65 ms. The cocktail-party noise was generated by combining a relatively dry recording from a cafeteria with the recordings of the male and female competing talkers mentioned earlier. The levels of the competing talkers were each 3 dB higher than that of the cafeteria noise. Prior to mixing, both competing talker recordings and the cafeteria noise were convolved with an impulse response from the Aachen Impulse Response database (office setting with dummy head at 1 m distance from the loudspeaker; Jeub *et al*., 2009) to introduce reverberation to the original recordings (*T*_60_ = 0.37 s). The spectra of the SSN and ICRA-5 noises, as well as that of the cafeteria noise used to create the cocktail-party noise, had been pre-processed to match the long-term average spectrum of the HINT speech material as closely as possible. The recording for the male and female competing talkers had not been spectrally shaped, as their spectra were deemed sufficiently close to the long-term average spectrum of the HINT material. The long-term average spectra and modulation spectra of the Danish HINT speech material and all maskers are presented in Figure 4.

**Figure 4.**
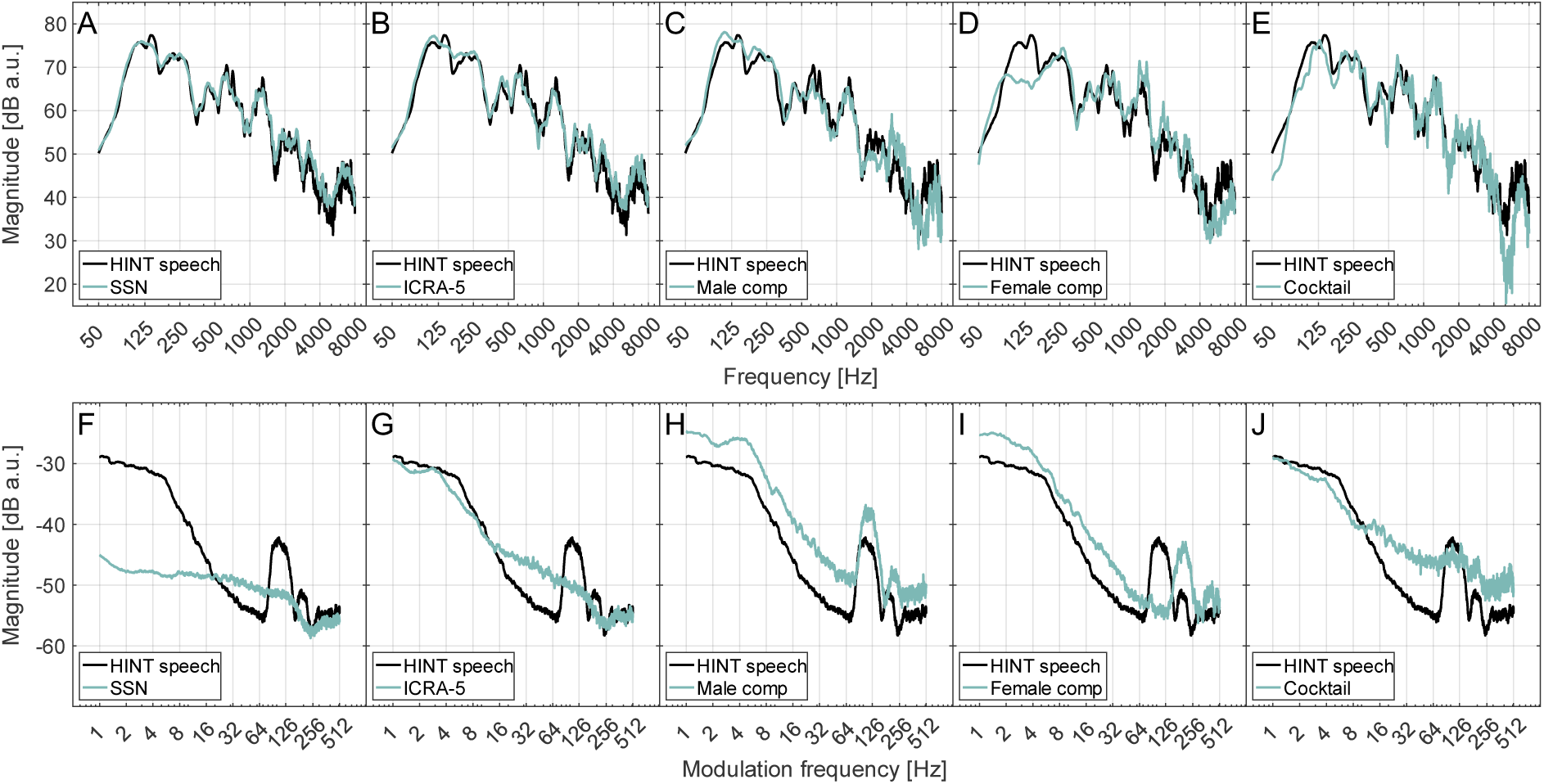
(Color online): Long-term average spectra (top row) and modulation spectra (bottom row) of the speech material and maskers. Each panel shows the spectra for the HINT speech material (black line) and one of the maskers (light blue line). Panels A and F: SSN; Panels B and G: ICRA-5; Panels C and H: Male competing talker; Panels D and I: Female competing talker; Panels E and J: Cocktail-party condition.

Each participant underwent training using all three training lists mixed with SSN. The rule defining the starting SNR (*SNR*_*start*_) for each masker and testing run was based on pilot data from two young NH listeners, whose SRTs were measured for all maskers. The pilot data indicated expected SRT offsets (*SRT*_*offset*_) relative to the SSN condition as follows: 0 dB for the male competing talker and cocktail-party conditions, −3 dB for the ICRA noise, and −9 dB for the female competing talker. Based on these offsets, *SNR*_*start*_for each masker was adjusted to be 6 dB below the expected SRTs, calculated relative to the SRT measured in the final training run with SSN (*SRT*_*final train*_) and rounded to the nearest integer (see Equation 1). Therefore, *SNR*_*start*_ was 6 dB lower than *SRT*_*final train*_ for the male competing talker, SSN, and cocktail-party conditions; 9 dB lower for the ICRA noise; and 15 dB lower for the female competing talker condition.

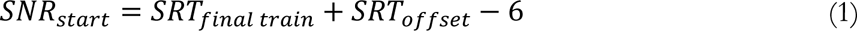

The SRT for each masking noise was determined by averaging two separate measurements using different lists, with masker presentation and lists order randomized across listeners. Both measurements for each masker were conducted consecutively.^4^ The SRTs were estimated using an adaptive 1-up 1-down procedure targeting 50%-correct performance with sentence-based scoring. A response was correct if all five words of the sentence were accurately repeated. The initial step size in SNR was 4 dB for the first four sentences, then reduced to 2 dB. A refinement rule was applied to the 5^th^ sentence: the change in SNR was computed as the average of the preceding four SNRs and the predicted next SNR, minus the immediately preceding SNR. The first sentence was repeated until the listener repeated it correctly. If the sentence required more than four repetitions, the run was terminated, and the starting SNR was increased by 2 dB before restarting. The SRT for a run was calculated as the average SNR from sentences 5 to 20, plus the SNR required for a fictitious 21^st^ sentence, determined by scoring the 20^th^ sentence (Soli and Wong, 2008). The scoring was performed by a trained native Danish speaker.

Target sentences were presented at a fixed level of 60 dB SPL, with masking noise looped continuously and adjusted in level to achieve the desired SNR. For the HI listeners, linear amplification was applied to both speech and masking noise using the Cambridge formula (CamEq; Moore and Glasberg, 1998), with a maximum gain of 30 dB at any frequency.^5^

### D. Statistical analysis

#### 1. Group-level analysis of SRTs

The degree of variability between the two SRT estimates obtained for each masker was assessed by calculating the standard deviation across the absolute differences in SRTs, *σ*_Δ|*SRT*|_. Notably, although small, the variability across all maskers was highest for the young NH listeners (*σ*_Δ|*SRT*|_= 2.43 dB) compared to the older NH listeners (*σ*_Δ|*SRT*|_= 1.76 dB) and older HI listeners (*σ*_Δ|*SRT*|_= 1.49 dB). Nevertheless, the degree of variability between SRT estimates was considered sufficiently small for all statistical analyses to be conducted using the SRTs averaged across both repetitions for each listener and masker. Analyses of variance (ANOVAs) of linear mixed-effects model fittings were performed to examine the differences between the listener groups. Two sets of analysis were conducted: First, the older NH group was compared to the young NH listeners to isolate the effects of age from hearing loss, and second, the older HI group was compared to the older NH listeners to isolate the effects of hearing loss from age-related effects. In both analyses, the listeners were treated as a random effect, while group classification and masker were considered as fixed effects.^6^ Post-hoc analyses were carried out using estimated marginal means, and a Holm-Bonferroni correction criterion was applied to account for multiple comparisons. The significance level for all analyses was set at 5%.

#### 2. Regression analysis

To assess whether a change in AM frequency selectivity contributes to the variance in SRTs, a multilinear regression model was built to relate age, audibility, working memory capacity, and AM frequency to SRTs across listeners. The SRTs from the condition showing the largest effect size across young and older NH listeners were chosen as the model’s outcome variable. The measure of age was taken as the listeners’ chronological age at the time of testing. The measure of audibility was taken as the 4-frequency pure-tone average threshold (PTA; where the audiometric thresholds at 0.5, 1, 2, and 4 kHz are averaged). The measure of working memory capacity was taken as the listeners’ RDS score. Finally, the measure of AM frequency selectivity was taken as the listeners’ DR_MTP_ at a 4-Hz target modulation frequency, shown in Figure 3. The model included all young and older NH listeners and 8 of the 9 older HI listeners, since a 4-Hz MTP could not be obtained by Regev *et al*. (2024a) for one of them. The multilinear regression model was built in steps, starting from a model including only age, by assessing whether adding the other predictors and their interactions significantly reduced the model’s residual sum of squares, using an F-test. The predictors were added in the above order (PTA, RDS, and DR_MTP_) and the addition of their interactions was only assessed if the main effect significantly improved the model. The significance level was set at 5%.

### III. RESULTS

#### A. Speech intelligibility

The SRTs for all listeners and groups are shown in Figure 5 for each masker. The young NH listeners are represented by squares, older NH listeners by triangles, and older HI listeners by circles. The statistical analysis comparing young and older NH listeners, focusing on the effects of age, revealed significant main effects of masker [*F_4,68_* = 274.2, *p* < 0.001] and listener group [*F_1,17_* = 17.56, *p* < 0.001], as well as a significant interaction between both factors [*F_4,68_* = 9.28, *p* < 0.001]. In the analysis comparing older NH and older HI listeners, examining the effects of hearing loss, significant main effects of masker [*F_4,64_* = 141.6, *p* < 0.001] and listener group [*F_1,16_* = 12.86, *p* = 0.002], as well as a significant interaction between both factors [*F_4,64_* = 5.32, *p* < 0.001], were observed. Post-hoc analyses revealed that the older NH listeners had significantly higher SRTs than the young NH listeners for the ICRA-5 condition [Effect size (*M*) = 1.39 dB, standard error (*SE*) = 0.68 dB], male competing talker [*M* = 4.53 dB, *SE* = 0.68 dB], female competing talker [*M* = 2.70 dB, *SE* = 0.68 dB], and cocktail-party condition [*M* = 1.53 dB, *SE* = 0.68 dB]. Additionally, the older NH listeners had significantly lower SRTs than the older HI listeners for the SSN condition [*M* = 2.36 dB, *SE* = 1.14 dB], ICRA-5 condition [*M* = 4.30 dB, *SE* = 1.14 dB], male competing talker [*M* = 3.60 dB, *SE* = 1.14 dB], and female competing talker [*M* = 5.52 dB, *SE* = 1.14 dB]. The increase in SRT for the older HI listeners in the cocktail-party condition approached, but did not reach, significance [*M* = 2.24 dB, *SE* = 1.14 dB, *p* = 0.06]. Furthermore, the post-hoc analyses revealed that the female competing talker resulted in significantly lower SRTs than all other maskers for all listener groups. The cocktail-party condition yielded the highest SRTs for the young NH listeners, while the male competing talker resulted in significantly higher SRTs than all other maskers for the older HI listeners. For the older NH listeners, both the male competing talker and cocktail-party conditions produced similar SRTs, which were significantly higher than those for all other maskers. Finally, the ICRA-5 noise resulted in significantly lower SRTs than the SSN for the young and older NH listeners, but not for the older HI listeners. Additional details and results from all post-hoc analyses can be found in Appendix A.

**Figure 5.**
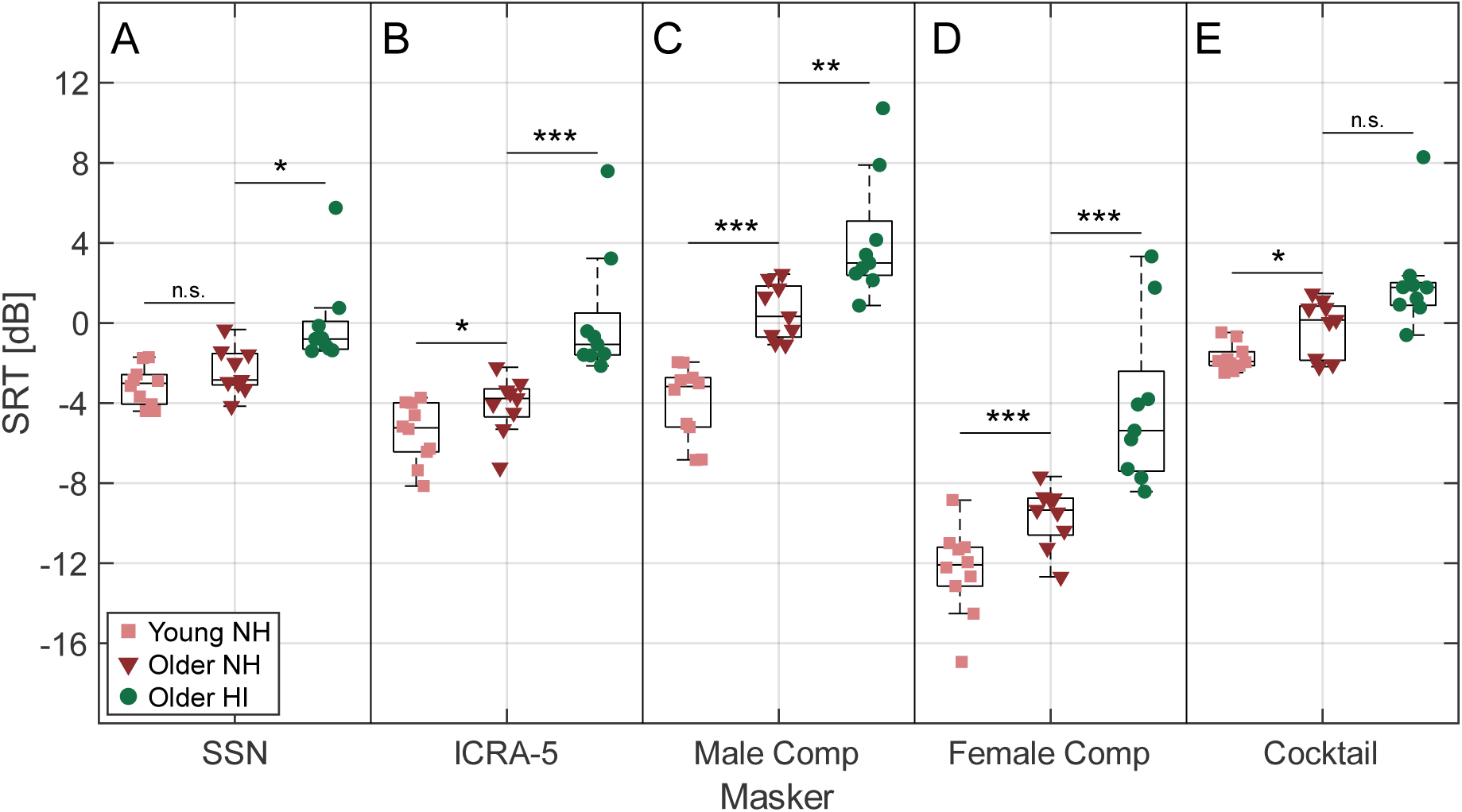
(Color online): Boxplots displaying the SRTs for all listeners. Each panel corresponds to a specific masker: Panel A: Speech-shaped noise (SSN), Panel B: ICRA-5 fluctuating noise, Panel C: Single male competing talker (same sex), Panel D: Single female competing talker (different sex), Panel E: Cocktail-party noise. The SRTs for the young NH listeners are represented by squares, older NH by triangles, and older HI by circles. The stars indicate the level of statistical significance of the differences between the groups separated by the horizontal black lines (*\* p < 0.05, ** p < 0.01, *** p < 0.001*), while “n.s.” denotes non-significant differences. Note that the young NH and older HI groups were not compared in the statistical analysis.

The multilinear regression analysis included the SRTs obtained with the male competing talker as the outcome variable, as that condition yielded the largest difference in SRTs between young and older NH listeners, as well as highest SRTs for both the older NH and HI listeners. The results showed that the best-fitting model included the main effects of age, PTA, and DR_MTP_, while RDS did not contribute to explaining the variance in SRTs. Adding DR_MTP_ to age and PTA [*F_1,23_* = 9.693, *p* = 0.005] significantly increased the amount of variance in SRT explained by the model (adjusted *R*^2^) by 8.4 percentage points to a total of 77%. No interaction term significantly increased the model’s *R*^2^. Figure 6 shows the scatter plots of the fitted SRTs as a function of the measured SRTs for each one of the models iteratively built (i.e., including age, age + PTA, or age + PTA + DR_MTP_ as predictors) in separate panels. The adjusted *R*^2^for each model is indicated in each panel. The best-fitting model’s regression coefficient (i.e., β weights) was larger for age (0.773) than for PTA and DR_MTP_ (0.303 and 0.350, respectively). The structure coefficients, i.e., the correlation of each predictor with the outcome variable estimated from all predictors in the model, showed that both age and PTA were positively correlated with the fitted SRTs, while DR_MTP_ was negatively correlated (0.896, 0.842, and −0.157, respectively).

**Figure 6:**
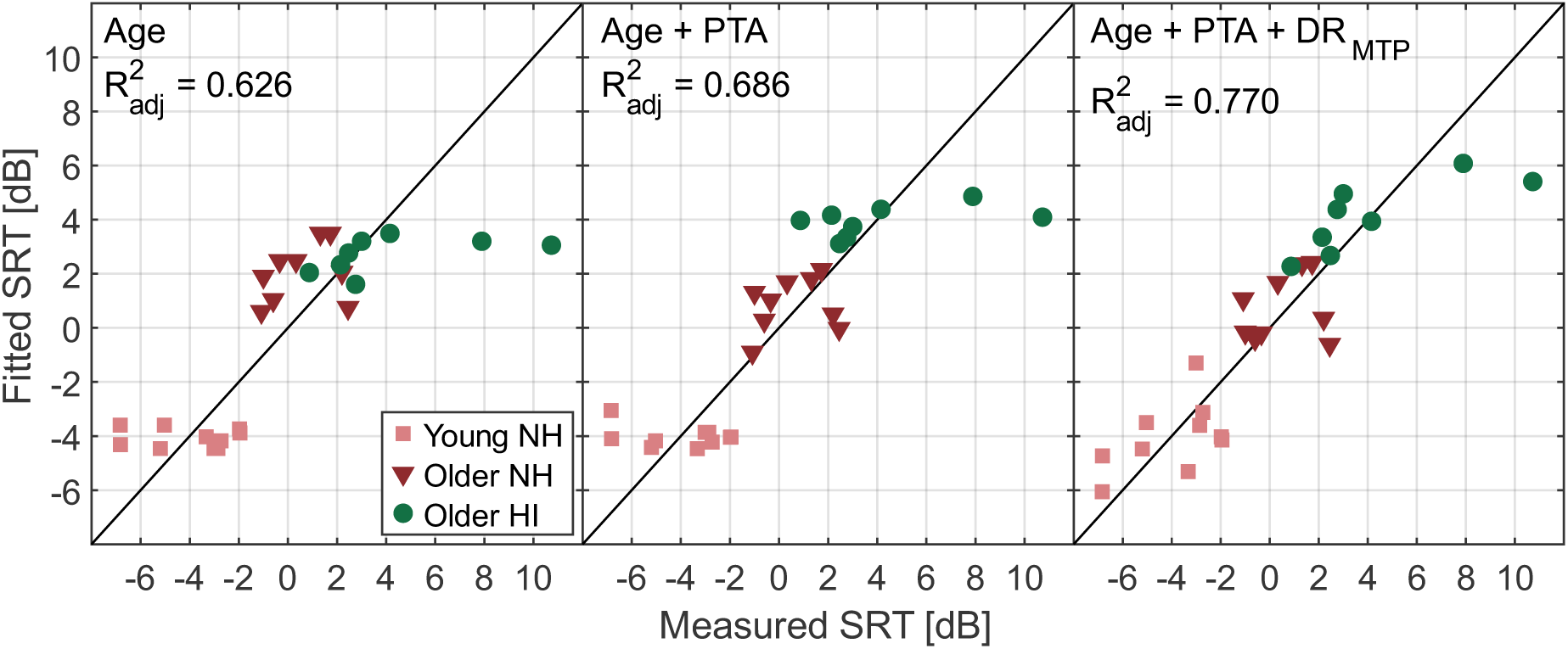
Scatter plot of the fitted SRTs as a function of the measured SRTs for each one of the regression models assessed. The predictors included in the model as well as the adjusted *R*^2^ are indicated in each panel. The SRTs for the young NH listeners are represented by squares, older NH by triangles, and older HI by circles. The diagonal line indicates identity. The models were iteratively built to include only age (left panel), age and PTA (middle panel), and finally age, PTA, and dynamic range of the 4-Hz MTP (DR_MTP_; right panel). The last model provided the best fit of the SRTs.

Since all predictors in the best-fitting model (age, PTA, and DR_MTP_) were likely somewhat collinear, a commonality analysis (Capraro and Capraro, 2001; Ray-Mukherjee *et al*., 2014) was implemented to assess the *R*^2^either attributable uniquely to each predictor or shared between them. The outcome showed that age had the greatest unique *R*^2^(22 %), while PTA and DR_MTP_ had smaller unique *R*^2^ (4.3 and 8.3 %, respectively). Furthermore, the commonality analysis revealed that age and PTA were highly collinear whereby they shared a large common *R*^2^ (48.8 %), and that age and DR_MTP_ were also somewhat collinear but shared a negative *R*^2^(−7.9%). The detailed results of the regression and commonality analyses can be found in Appendix B.

## IV. DISCUSSION

The aim of the present study was to investigate whether the findings from speech intelligibility tests would be consistent with the AM frequency selectivity data collected in previous studies by Regev *et al*. (2023, 2024a). These previous investigations demonstrated an age-related decline in AM frequency selectivity and an effective sharpening of AM tuning with hearing loss (see Figure 2 and Figure 3). However, the latter was proposed to be an “apparent” result of reduced cochlear compression (Regev *et al*., 2024a), rather than a true sharpening of the theoretical modulation filters. Thus, older HI listeners are also hypothesized to experience an age-related decline in AM selectivity. A subset of the listener cohort measured by Regev *et al*. (2023, 2024a) was considered here. It was expected that the age-related reduction in AM frequency selectivity would contribute to poorer speech intelligibility in older NH and HI listeners compared to young NH listeners.

The results revealed an overall decline in speech intelligibility with age, which was further exacerbated by the presence of hearing loss. These findings are consistent with previous studies that have shown a deterioration of speech intelligibility with age (e.g., Rajan and Cainer, 2008; Helfer and Vargo, 2009; Schoof and Rosen, 2014; Füllgrabe *et al*., 2015; Gordon-Salant and Cole, 2016; Goossens *et al*., 2017; Decruy *et al*., 2019) and hearing loss (e.g., Glasberg and Moore, 1989; Humes and Christopherson, 1991; Frisina and Frisina, 1997; see Humes and Dubno, 2010 for a review). Additionally, a multilinear regression model relating the SRTs obtained with a male competing talker to age, PTA, and DR_MTP_ (a measure of AM frequency selectivity) showed that DR_MTP_ significantly contributed to explaining the variance in SRTs across all listeners, increasing the model’s *R*^2^ by 8 percentage points.

The observed increase in SRTs with age aligns with the age-related reduction in AM frequency selectivity reported by Regev *et al*. (2023). Broader modulation filters, indicative of reduced AM selectivity, may lead to increased AM masking, resulting in a decrease in the SNR_env_ at the output of these filters and potentially poorer speech intelligibility in noise. However, the lack of a significant age-related increase in SRT for the SSN condition is inconsistent with this notion. Since the SSN contains intrinsic envelope fluctuations that act as a relatively broadband modulation masker (as argued by Stone *et al*., 2012), one might have expected the broader AM tuning found in older NH listeners to lead to an increase in AM masking. However, the SRTs obtained with the SSN likely reflect aspects of EM rather than AM masking. This is consistent with the extensive overlap observed in the audio spectra of the SSN and HINT speech material (Figure 4, panel A), compared to the highly positive SNR_env_ observed when comparing their modulation spectra at low modulation frequencies (Figure 4, panel F). These findings indicate that the SSN provided more EM than AM masking. Conversely, the differences in modulation spectra between the target speech and masking noise were much smaller for the other maskers, suggesting that AM masking (and thus potentially AM frequency selectivity) may have played a more prominent role in those cases (Figure 4, panels G-J). Notably, the older NH listeners exhibited a reduced release from masking when exposed to the fluctuating ICRA-5 noise compared to the young NH listeners, consistent with previous studies (Dubno *et al*., 2002, 2003; Grose *et al*., 2009; Goossens *et al*., 2017). In addition to potentially being the result of an age-related increase in forward masking (e.g., Gifford *et al*., 2007), the reduced release from masking observed in the older NH listeners may also be attributed to increased AM masking due to the loss of AM frequency selectivity, which may have limited the benefit derived from the slow temporal fluctuations in the ICRA-5 condition.

Regarding the effects of hearing loss, the higher SRTs observed in the older HI listeners compared to the older NH group are consistent with the expectation that the increase in AM frequency selectivity observed in HI listeners (as reported in Regev *et al*., 2024a) would *not* lead to improved speech intelligibility. Instead, it was hypothesized that older HI listeners also experience an age-related loss of AM selectivity that contributes to poorer speech intelligibility. This was based on the proposition by Regev *et al*. (2024a) that the greater AM frequency selectivity observed with hearing loss is an apparent effect resulting from improved detection of the AM targets during temporal dips in the masker, due to reduced cochlear compression. Since the loss of compression is also associated with other supra-threshold deficits, such as loudness recruitment, poorer recovery from forward masking, and reduced frequency selectivity (Glasberg and Moore, 1986, 1989; Oxenham and Moore, 1997; Moore and Oxenham, 1998), it is evident that the process potentially underlying the apparent improved AM frequency selectivity is also linked to the mechanisms underlying poorer speech intelligibility in HI listeners. However, while poorer speech intelligibility was expected in the HI listeners, disentangling the potential impact of the effective sharpening in AM tuning with hearing loss on speech intelligibility is not straightforward. It remains unclear whether the process underlying the apparent sharpening of AM tuning with hearing loss a) improves SRTs but is outweighed by co-occurring supra-threshold deficits typically associated with hearing loss, or b) itself ultimately impairs speech intelligibility. The group-level results obtained for the older HI listeners do not clarify whether an age-related reduction in AM selectivity contributed to the increase in SRTs, beyond the effects of hearing loss.

However, the results of the multilinear regression analysis allowed for a separate investigation of these effects, further supporting the existence of a link between alterations in AM frequency selectivity and speech intelligibility challenges in noise for older listeners. The addition of DR_MTP_ to the model significantly increased the amount of variance in SRTs explained by the model. Furthermore, the negative structure coefficient for DR_MTP_ indicated that an increase in the SRTs estimated by the best-fitting model (which included age, PTA, and DR_MTP_) is linked with a decrease in DR_MTP_. This finding aligned with the expectation that poorer AM frequency selectivity (quantified by a lower DR_MTP_) would be linked with higher SRTs.^7^ This link, identified in a model including all listeners, is also broadly consistent with the proposition by Regev *et al*. (2024a) that older HI listeners experience an age-related reduction in AM frequency selectivity, which is concealed in the MTPs by a perceptual benefit stemming from the loss of cochlear compression (represented by PTA in the regression model).

Employed to address the likely collinearity across predictors, the commonality analysis revealed that DR_MTP_ acts as a suppressor, as shown by the negative *R*^2^ shared between DR_MTP_ and age (Capraro and Capraro, 2001). As such, the addition of DR_MTP_ to the model helped increase the *R*^2^uniquely attributable to age, thus increasing the model’s total *R*^2^. Within the model, age can be seen as a “noisy term” representing the sum of all factors underlying auditory and cognitive processing deficits in older listeners that are distinct from the effects of peripheral hearing loss and reduced working-memory capacity. Such factors, which are likely somewhat collinear, may include cochlear synaptopathy (Parthasarathy *et al*., 2019a), reduced neural inhibition (Caspary *et al*., 2008), or poorer temporal envelope processing (Anderson and Karawani, 2020) such as a loss of AM frequency selectivity (Regev *et al*., 2023). Hence, the greater *R*^2^resulting from the explicit addition of a measure of AM frequency selectivity to the regression model can reasonably be interpreted as resulting from a “denoising” of the age term, allowing it to more accurately account for the variance in SRTs.

However, it should be noted that the significant contribution of DR_MTP_ to the regression model was not observed when the model was applied to SRTs for maskers other than the male competing talker or DR_MTP_ derived from MTPs at modulation rates other than 4 Hz. For the SSN and cocktail-party conditions, this may partly result from the limited variance in SRTs across listeners (see Figure 5 A and E). For the ICRA-5 and female competing talker conditions, only the main effect of PTA significantly contributed to explaining the variance in SRTs. This dominance of the PTA effect and the unexpected lack of an age effect may arise from the overlap in SRTs between young and older NH listeners (though significantly different) for these conditions, while the older HI listeners exhibited substantially higher SRTs compared to the NH groups (see Figure 5 B and D). Hence, the age effect in these conditions may not have been strong enough for an age-related loss in AM frequency selectivity to influence intelligibility. Overall, while the reasons for the inconsistent effects of age and DR_MTP_ remain unclear, the specific selection of the SRTs for the male competing talker and the DR_MTP_ derived from a 4-Hz modulation rate was intentional in order to maximize the variability of dependent and independent variables in the regression model. The male competing talker yielded the largest SRT differences between young and older NH listeners and the highest SRTs for the older NH and HI listeners compared to other maskers. Additionally, the 4-Hz MTP showed the largest differences in AM selectivity between young and older NH listeners (Regev *et al*., 2023). Finally, the 4-Hz MTP likely reflects the prominent role of this modulation rate in speech perception (e.g., Elliott and Theunissen, 2009; Varnet *et al*., 2017), making it more perceptually relevant than MTPs obtained at other rates.

Notably, the present findings indicate that declines in speech intelligibility with age and hearing loss are most pronounced in contexts involving high cognitive demand and informational masking (IM), such as in the presence of competing-talker maskers. IM includes factors that complicate the separation target and masker speech signals, such as stream segregation (Brungart et al., 2001; Darwin et al., 2003; Schneider et al., 2007; Kidd et al., 2016; see Kidd et al., 2008 for a review)). In this study, both older NH and older HI listeners exhibited lower RDS scores compared to the young NH listeners (significantly and marginally so, respectively). Such reduced cognitive abilities have been implicated in age-related increases of SRTs under conditions of IM (Rajan and Cainer, 2008; Gordon-Salant and Cole, 2016; Goossens *et al*., 2017) and in the pronounced effects of hearing loss in cognitively demanding settings (e.g., Schneider *et al*., 2010; Goossens *et al*., 2017). Thus, the poorer speech intelligibility observed in the older NH and HI listeners in the present study may partly reflect reduced working-memory capacity. However, significant effects of hearing loss and age were also found for the ICRA-5 condition, which does not impose high IM or cognitive demand. Furthermore, the multilinear regression analysis showed that RDS scores did not contribute to explaining the variance in SRTs for the male competing talker across all listeners, suggesting that working-memory capacity did not significantly influence the observed speech intelligibility results. It is important to note that working-memory capacity is only one of several cognitive abilities that may be reduced in older listeners, potentially contributing to challenges in noisy environments (e.g., Dryden *et al*., 2017). Overall, while reduced cognitive ability may have partly mediated the effects of age and hearing loss on speech intelligibility, the results also point to supra-threshold deficits. These may involve age-related temporal processing deficits, as suggested by perceptual (Pichora-Fuller and Souza, 2003; Pichora-Fuller and MacDonald, 2008; Helfer and Vargo, 2009; Füllgrabe et al., 2015; see Fitzgibbons and Gordon-Salant, 2010 for a review) and neurophysiological research (e.g., Anderson et al., 2012; Decruy et al., 2019; Parthasarathy et al., 2019b; Roque et al., 2019; McFarlane and Sanchez, 2024; for reviews see Parthasarathy et al., 2019a and Anderson and Karawani, 2020), or reduced cochlear compression (e.g., Glasberg and Moore, 1989; Oxenham and Moore, 1997; Moore and Oxenham, 1998; Moore, 2003). The hypothesis that an age-related reduction in AM frequency selectivity contributes to increased SRTs aligns with these findings from previous studies.

Although the data from the present study revealed significant group-level and regression results supporting a relationship between changes in AM tuning and speech intelligibility in noise for older NH and HI listeners, it is important to consider several co-occurring factors that may have influenced the findings. Even though all the older NH listeners exhibited clinically normal hearing up to 4 kHz, their audiometric thresholds were on average 8 dB higher than those of the young NH listeners between 125 Hz and 8 kHz, with the greatest differences occurring at 8 kHz. Although these differences in audiometric thresholds are small, previous studies have suggested that they can have a considerable effect on speech intelligibility (Dubno and Ahlstrom, 1997; Dubno *et al*., 2000), possibly due to related supra-threshold processing deficits (e.g., Léger *et al*., 2012). Moreover, the older NH listeners included in the present study may have experienced reduced audibility at extended high frequencies (> 8 kHz), which have also been suggested to play a role in speech intelligibility (e.g., Hunter *et al*., 2020; Polspoel *et al*., 2022). These aspects may have contributed to the observed age-related increase in SRTs and are consistent with the significant effect of PTA found in the regression analysis of the SRTs.

Concerning the experimental approach, since the tests provided listeners with cues beyond the temporal envelope, factors such as IM, EM, and stream segregation likely interacted with age- and hearing loss-related perception deficits, as well as the age-related reduction in working-memory capacity discussed above. Therefore, the connection between SRTs and AM frequency selectivity suggested by the present results warrants further investigation. Notably, a regression analysis using the SRTs from the ICRA-5 condition, which is not affected by IM, revealed that only PTA significantly contributed to explaining the variance in SRTs. This suggests that the effect of age on SRTs in this condition was not sufficiently strong for an age-related loss of AM selectivity to play a role. Overall, an experimental paradigm limiting the perceptual cues available in the speech intelligibility test to the temporal envelope could help establish a more definitive link between declines in AM selectivity and speech intelligibility.

Finally, the outcomes of the multilinear regression model are limited by the bimodal distributions of age and PTA across listeners, which may have contributed to the strong effect of age found in the model. Since the studies by Regev *et al*. (2023, 2024a) focused on groups of young NH, older NH, and older HI listeners, both the age and PTA dimensions are constrained to low or high values. A study recruiting a larger sample with more continuous distributions of age and PTA would enable a stronger regression model and more conclusive outcomes. Moreover, recruiting NH listeners across a continuum of ages would allow for a regression analysis focusing exclusively on the effects of AM selectivity on SRTs in isolation from hearing loss, potentially providing clearer evidence of a link between these measures.

## V. SUMMARY AND CONCLUSION

Speech intelligibility in noise was examined in three groups: young listeners with normal hearing (aged 22-28 years), older listeners with normal hearing (aged 57-77 years), and older listeners with hearing impairment (aged 64 – 77) years. All participants had previously taken part in AM masking studies investigating AM frequency selectivity. These studies revealed an age-related reduction in AM tuning (Regev *et al*., 2023), and an apparent sharpening in AM tuning with hearing loss (Regev *et al*., 2024a), which was attributed to reduced cochlear compression rather than a true sharpening of the theoretical modulation filters. This study aimed to determine whether the effects of age and hearing loss on speech intelligibility in noise support the hypothesis that an age-related reduction in AM tuning in older NH and HI listeners negatively impacts speech intelligibility.

Consistent with the hypothesis, the results showed an age-related decline and an additional negative effect of hearing loss on speech intelligibility, especially for fluctuating maskers. Furthermore, a multilinear regression analysis showed that adding a measure of AM frequency selectivity to a model using age and audibility as predictors significantly increased the amount of variance in SRTs explained across all listeners. This finding supported the assumption that an age-related loss of AM frequency selectivity is linked to speech intelligibility difficulties in noise in older NH and HI listeners, once audibility is accounted for. Moreover, the results broadly aligned with the proposition made by Regev *et al*. (2024a) that the improved AM tuning in listeners with hearing impairment is an “apparent” effect resulting from reduced peripheral compression, which is known to contribute to other supra-threshold deficits that impair speech intelligibility.

Overall, this study’s results support the notion that an age-related reduction in AM frequency selectivity may contribute to speech intelligibility challenges in noise and warrant further investigation into this potential mechanism. To more directly assess the effects of AM tuning, and a loss thereof, on speech intelligibility, future studies may focus on speech-intelligibility testing paradigms that specifically examine envelope-based cues.

## ACKNOWLEDGMENTS

The authors thank Borgný Súsonnudóttir Hansen for her contribution to the data collection. The authors also thank Christian Stender Simonsen for kindly sharing the experimental framework used to run the listening test, as well as several of the signal recordings, and Jens Hjortkjær and Jonatan Märcher-Rørsted for kindly providing their implementations of the reverse digit span test and of the Cambridge formula (CamEq; Moore and Glasberg, 1998). We thank Dr. Christopher Brown and two anonymous reviewers for their insightful comments on earlier versions of this manuscript. Finally, the authors thank Raul Sanchez-Lopez for helpful insights and discussions. This study was supported by the Oticon Centre of Excellence for Hearing and Speech Sciences (CHeSS).

## AUTHOR DECLARATIONS

### Ethics Approval

All participants provided written informed consent and were offered financial compensation for their participation. Ethical approval for the study was provided by the Science Ethics Committee of the Capital Region in Denmark (reference H-16036391).

## DATA AVAILABILITY

This manuscript has been submitted to the Journal of the Acoustical Society of America. Once it has been accepted, the data that support the findings of this study will be openly available from: Regev, J., Zaar, J., Relaño-Iborra, H., and Dau, T. (2024). “Dataset for: ‘Investigating the effects of age and hearing loss on speech intelligibility and amplitude modulation frequency selectivity’”, Technical University of Denmark. Dataset. https://doi.org/10.11583/DTU.25771884.

### APPENDIX

#### A. Post-hoc analyses

Table 1 summarizes the results of the post-hoc analyses that compared the SRTs obtained with young and older NH listeners for each masker.

**Table 1:** Post-hoc analysis of the differences in SRT between young and older NH listeners for each masker. The effect sizes (M), standard errors (SE, and corrected p-values are reported. The significant p-values are highlighted in bold.

| Masker | M [dB] | SE [dB] | Corrected p-value |
| --- | --- | --- | --- |
| SSN | 0.75 | 0.68 | 0.279 |
| ICRA-5 | 1.39 |  | **0.047** |
| Male competing | 4.53 |  | **< 0.001** |
| Female competing | 2.70 |  | **< 0.001** |
| Cocktail party | 1.53 |  | **0.030** |

Table 2 summarizes the results of the post-hoc analyses that compared the SRTs obtained with older NH and older HI listeners for each masker.

**Table 2:**
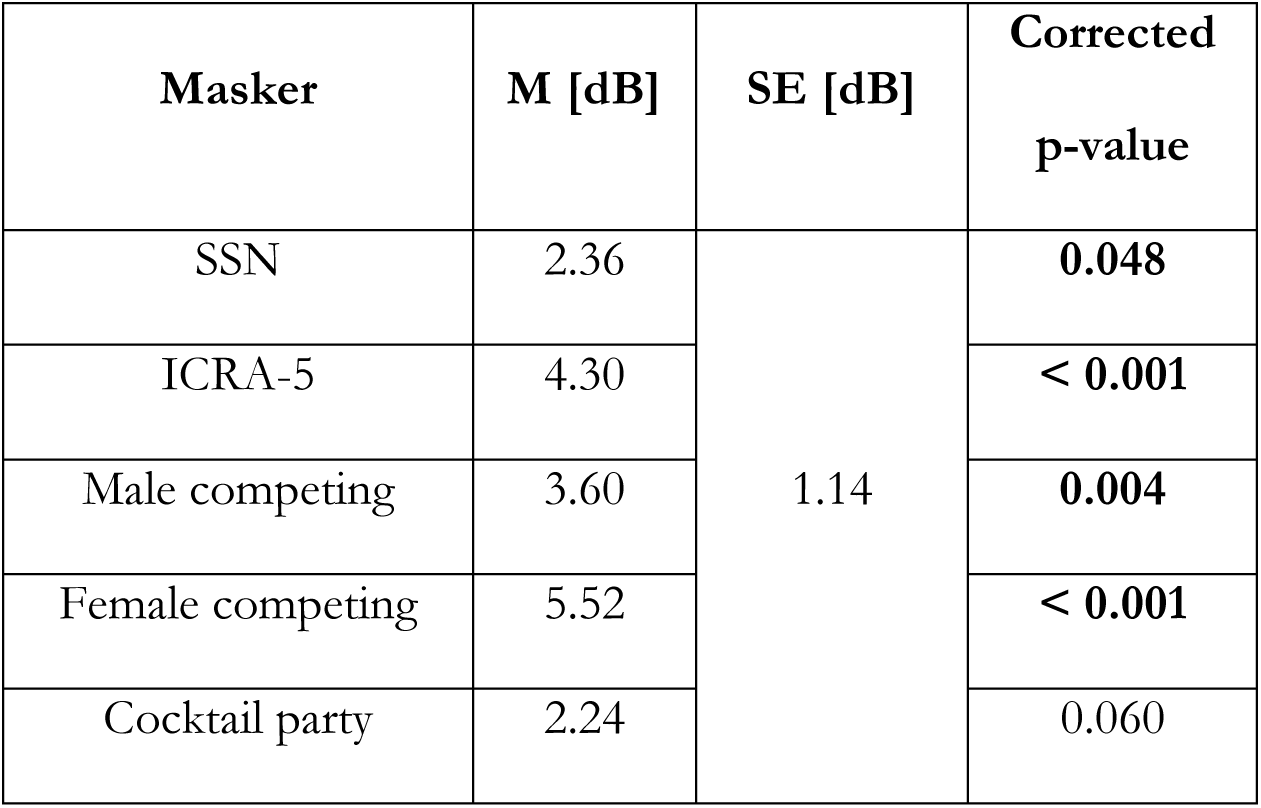
Post-hoc analysis of the differences in SRT between older NH and older HI listeners for each masker. The effect sizes (M), standard errors (SE, and corrected p-values are reported. The significant p-values are highlighted in bold.

| Masker | M [dB] | SE [dB] | Corrected p-value |
| --- | --- | --- | --- |
| SSN | 2.36 | 1.14 | **0.048** |
| ICRA-5 | 4.30 |  | **< 0.001** |
| Male competing | 3.60 |  | **0.004** |
| Female competing | 5.52 |  | **< 0.001** |
| Cocktail party | 2.24 |  | 0.060 |

Table 3 summarizes the results of the post-hoc analyses that compared the SRTs obtained with the different maskers for each of the three listener groups (young NH, older NH, and older HI). Since this post-hoc analysis was run twice for the older NH listeners (once for each statistical model applied), the values reported reflect the most conservative *p*-values and standard errors, obtained from the model fitted to the older NH and older HI listeners. However, the effect sizes and overall statistical outcomes did not differ between the analyses run in the two models.

**Table 3:**
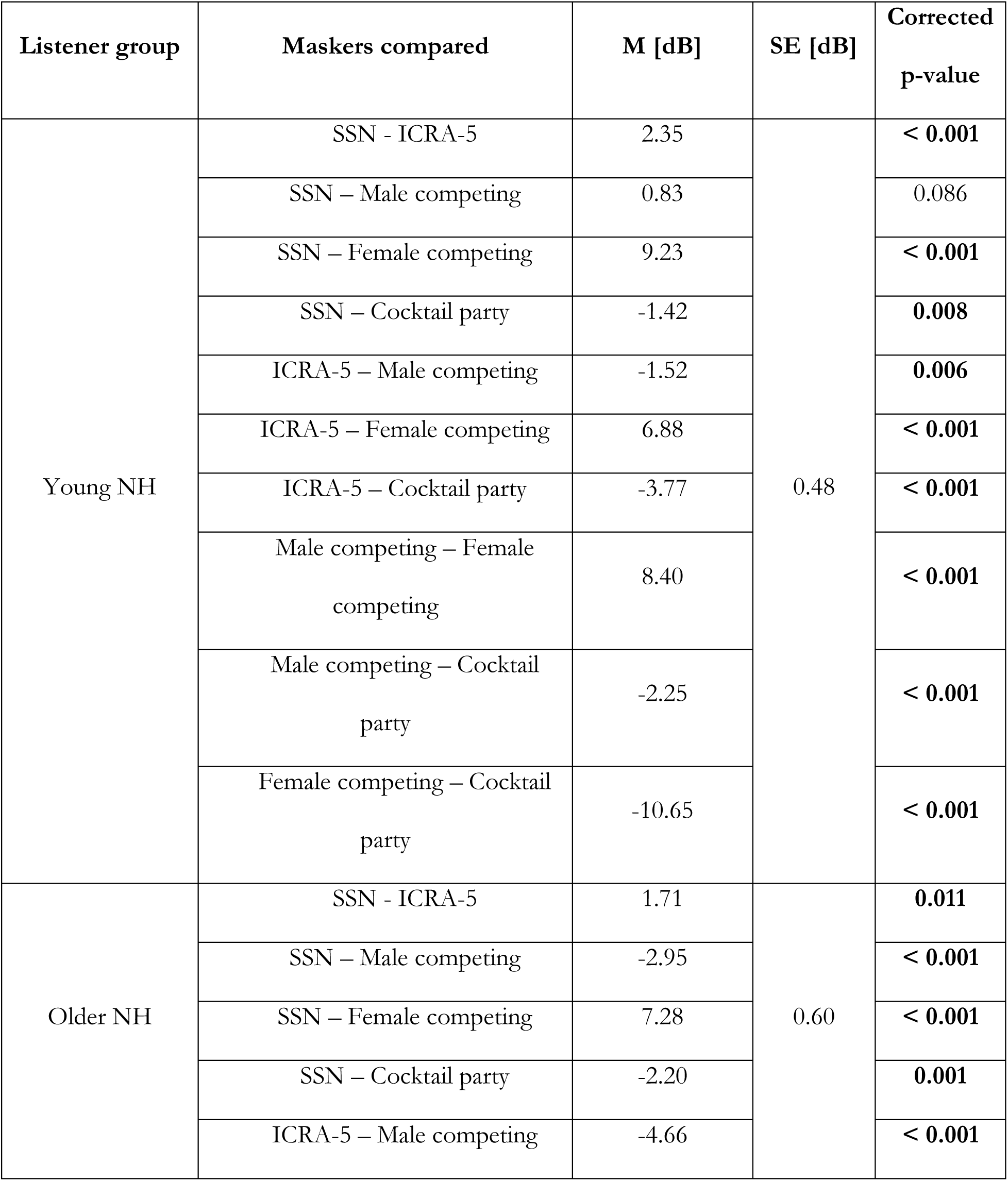
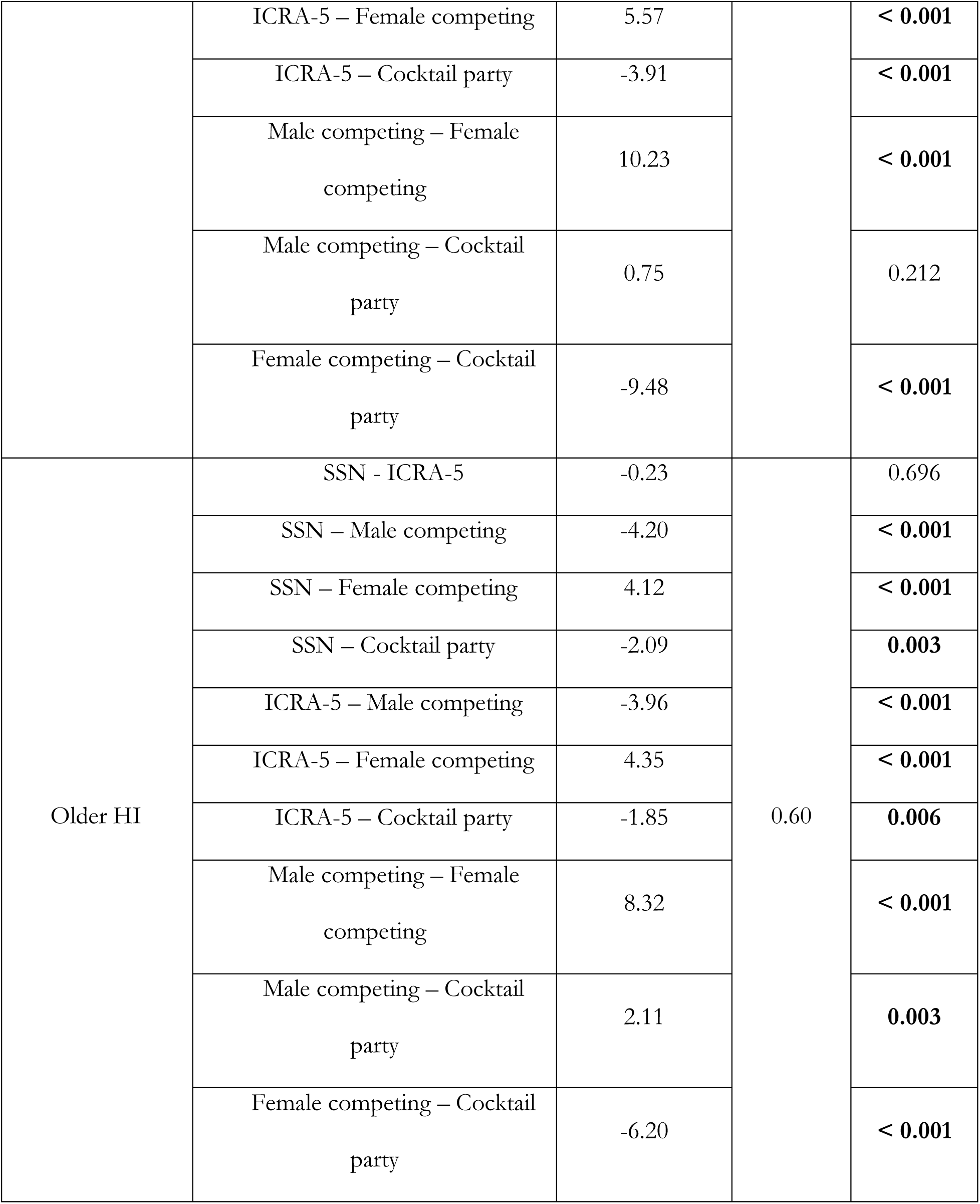
Post-hoc analysis of the differences in SRT between older the different maskers for each group. The effect sizes (M), standard errors (SE, and corrected p-values are reported. The significant p-values are highlighted in bold.

| Listener group | Maskers compared | M [dB] | SE [dB] | Corrected p-value |
| --- | --- | --- | --- | --- |
| Young NH | SSN - ICRA-5 | 2.35 | 0.48 | **< 0.001** |
|  | SSN – Male competing | 0.83 |  | 0.086 |
|  | SSN – Female competing | 9.23 |  | **< 0.001** |
|  | SSN – Cocktail party | -1.42 |  | **0.008** |
|  | ICRA-5 – Male competing | -1.52 |  | **0.006** |
|  | ICRA-5 – Female competing | 6.88 |  | **< 0.001** |
|  | ICRA-5 – Cocktail party | -3.77 |  | **< 0.001** |
|  | Male competing – Female competing | 8.40 |  | **< 0.001** |
|  | Male competing – Cocktail party | -2.25 |  | **< 0.001** |
|  | Female competing – Cocktail party | -10.65 |  | **< 0.001** |
| Older NH | SSN - ICRA-5 | 1.71 | 0.60 | **0.011** |
|  | SSN – Male competing | -2.95 |  | **< 0.001** |
|  | SSN – Female competing | 7.28 |  | **< 0.001** |
|  | SSN – Cocktail party | -2.20 |  | **0.001** |
|  | ICRA-5 – Male competing | -4.66 |  | **< 0.001** |
|  | ICRA-5 – Female competing | 5.57 |  | **< 0.001** |
|  | ICRA-5 – Cocktail party | -3.91 |  | **< 0.001** |
|  | Male competing – Female competing | 10.23 |  | **< 0.001** |
|  | Male competing – Cocktail party | 0.75 |  | 0.212 |
|  | Female competing – Cocktail party | -9.48 |  | **< 0.001** |
| Older HI | SSN - ICRA-5 | -0.23 | 0.60 | 0.696 |
|  | SSN – Male competing | -4.20 |  | **< 0.001** |
|  | SSN – Female competing | 4.12 |  | **< 0.001** |
|  | SSN – Cocktail party | -2.09 |  | **0.003** |
|  | ICRA-5 – Male competing | -3.96 |  | **< 0.001** |
|  | ICRA-5 – Female competing | 4.35 |  | **< 0.001** |
|  | ICRA-5 – Cocktail party | -1.85 |  | **0.006** |
|  | Male competing – Female competing | 8.32 |  | **< 0.001** |
|  | Male competing – Cocktail party | 2.11 |  | **0.003** |
|  | Female competing – Cocktail party | -6.20 |  | **< 0.001** |

#### B. Regression analysis

Table 4 summarizes the details of the best-fitting model used in the multilinear regression analysis.

**Table 4:**
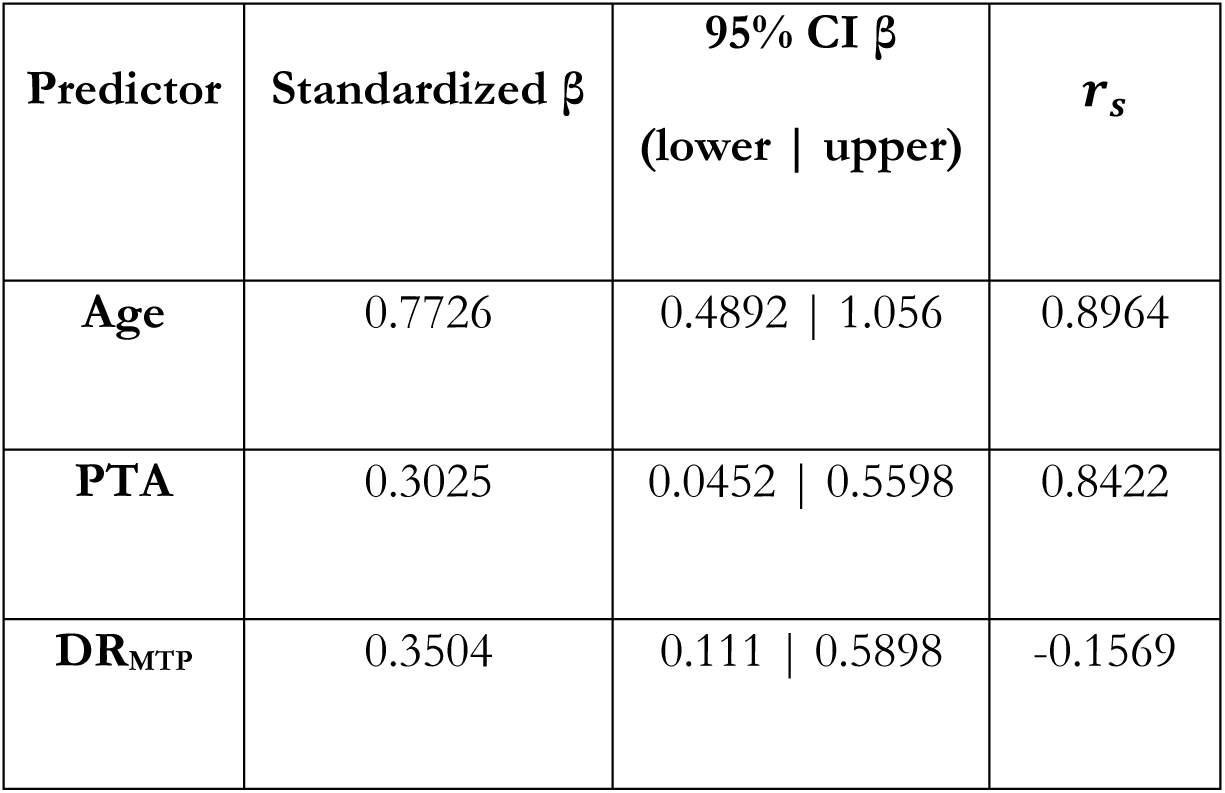
Details of the multilinear regression analysis relating the SRTs obtained for the male competing talker to age, PTA, and DR_MTP_. For each predictor, the table reports the standardized regression coefficients (β), the 95%-confidence interval for β (lower | upper), as well as the structure coefficients (*r*_*s*_).

| Predictor | Standardized $\beta$ | 95% CI $\beta$<br>(lower \| upper) | $r_s$ |
| --- | --- | --- | --- |
| Age | 0.7726 | 0.4892 \| 1.056 | 0.8964 |
| PTA | 0.3025 | 0.0452 \| 0.5598 | 0.8422 |
| $DR_{MTP}$ | 0.3504 | 0.111 \| 0.5898 | -0.1569 |

Table 5 summarizes the details of the commonality analysis conducted on the best-fitting model (relating SRTs to age, PTA, and DR_MTP_), which identifies the *R*^2^ either uniquely attributable to each predictor or shared among them.

**Table 5:** Outcome of the commonality analysis applied on the best-fitting model, that related SRTs obtained for a male competing talker to age, PTA, and the 4-Hz MTP’s average dynamic range (DR_MTP_). The table reports the amount of variance in SRTs explained (*R*^2^) either uniquely explained by each predictor (single variables) or commonly shared between predictors (two or more variables).

| Predictor | Age | PTA | $DR_{MTP}$ | Age and PTA | Age and $DR_{MTP}$ | PTA and $DR_{MTP}$ | Age, PTA, and $DR_{MTP}$ | Total |
| --- | --- | --- | --- | --- | --- | --- | --- | --- |
| $R^2$ [%] | 21.96 | 4.33 | 8.30 | 48.78 | -7.89 | 2.49 | -1.00 | 76.97 |

<sup>2</sup> Except for one listener who showed a gap of 20 dB at 1 kHz.

## Footnotes

1 The classification of the audiograms was performed using the lowest mean squared error between the standard audiograms and the collected audiometric thresholds at frequencies from 1 to 8 kHz.

2 Except for one listener who showed a gap of 20 dB at 1 kHz.

3 This was shown by the post-hoc analysis run on a linear model ANOVA using estimated marginal means. The older NH listeners had significantly lower RDS scores than the young NH and older HI listeners [*M* = 0.21, *SE* = 0.05 dB; *M* = 0.13, *SE* = 0.05 dB, respectively].

4 Due to a mistake in the audibility compensation, one of the older HI listeners was tested twice on the entire HINT corpus. However, more than 7 weeks separated the two testing sessions. Nielsen and Dau (2011) reported an average improvement in SRT of only 0.5 dB across two sessions separated by three weeks. Hence, the current case was likely not affected by memorization of the speech material.

5 The amplification produced aided thresholds of at most 35 dB until 4 kHz, 45 dB at 6 kHz, and 55 dB at 8 kHz. Hence, while reasonable audibility was achieved at low frequencies, audibility was not fully restored to levels comparable to those of the older NH listeners. However, a maximum gain of 30 dB was applied to ensure that no listener was exposed to uncomfortable or potentially hazardous sound levels during the speech intelligibility test, where the unaided masker levels could reach up to 80 dB SPL.

6 Levene’s test was used to assess the homoskedasticity of the residuals and Shapiro-Wilk’s test was used to assess the normality of residual’s distribution. Shapiro-Wilk’s test was not passed for either model, likely due to the relatively small group sizes and to the presence of outliers in the older HI data. However, no other aspect of the data justified the removal of the outliers, and they were therefore kept in the dataset. Furthermore, the average fittings of the models to the experimental data were found to be accurate, and the models were hence considered as appropriate.

7 Structure coefficients are employed here since they have been suggested to be more robust than regression coefficients in cases involving multicollinearity among predictors (Thompson and Borrello, 1985; Kraha *et al*., 2012)

