## Supplementary Figure 1 for "Investigating the effects of age and hearing loss on speech intelligibility and amplitude modulation frequency selectivity"

**Illustration of the derivation of the MTP's average dynamic range** $(DR_{MTP})$

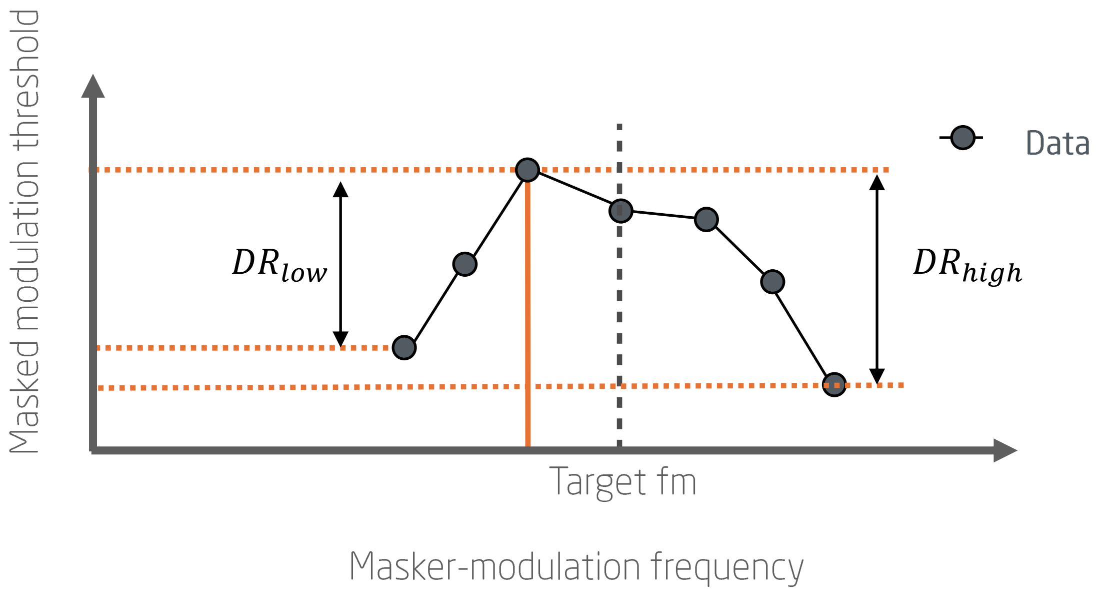


If $DR_{low} > 0$ and $DR_{high} > 0$

$$DR_{MTP} = \frac{DR_{low} + DR_{high}}{2}$$

If $DR_{low} = 0$

$$DR_{MTP} = DR_{high}$$

If $DR_{high} = 0$

$$DR_{MTP} = DR_{low}$$

Supplementary Figure 1: Illustration of the derivation of the MTP's average dynamic range ($DR_{MTP}$), obtained by averaging the dynamic ranges on the low and high skirts of the MTP ($DR_{low}$ and $DR_{high}$, respectively) around the MTP's peak. If the peak is located at either extreme, the $DR_{MTP}$ is taken as the dynamic range across the whole pattern.
